# 3D printing and deep learning enable holistic and dynamic analyses of tens of thousands of parasites infecting hundreds of genotypes

**DOI:** 10.64898/2026.08.13.744426

**Authors:** Siyuan Wei, Jie Zhou, Olaf Prosper Kranse, Unnati Sonawala, Gang Sun, Ziyang He, Beatrice Senatori, Clement Pellegrin, Andrea Díaz-Tendero Bravo, Roberta Healey, Victor Hugo Moura de Souza, Vincent C.T. Hanlon, George Harpum, Tithira Wijayathilake, Adela Gaja Jezierska-Suwinska, Anika Damm, Kerry VerMeulen, Thomas Baum, Lida Derevnina, Ji Zhou, Sebastian Eves-van den Akker

**Affiliations:** Department of Plant Sciences, University of Cambridge, Cambridge, CB3 0LE, UK; Academy for Advanced Interdisciplinary Studies, Nanjing Agricultural University, Nanjing, 210095, China; Department of Plant Pathology, Entomology and Microbiology, Iowa State University, Ames, IA 50011, USA; Cambridge Crop Research, National Institute of Agricultural Botany (NIAB), Cambridge, CB3 0LE, UK; University of the Chinese Academy of Sciences, Beijing 100049, China

## Abstract

Host-parasite interactions are dynamic systems, where parasites usually outnumber hosts by one or more orders of magnitude. However, our understanding is often limited by the assessment of parts of the host, at arbitrary time points, and/or aggregate parasite responses. Here we combined custom-built 3D-printed hardware and deep-learning-based algorithms to enable holistic (i.e. all infecting individuals on the whole plant), spatio-temporal, and parasite-centric analyses of plant-parasitism by nematodes from timelapse videos of infection over months. In so doing, we tracked the dynamic growth and development of all individual parasites, at the organismal level, for thousands of hosts across hundreds of genotypes of *Arabidopsis thaliana*. Categorising traits into the static (i.e. in an acquired image at a given time point) and dynamic (i.e. phenotypic changes over time), we revealed a greater extent of host-genetic control of parasite traits, and new physiological limits of the species under these conditions. Using this capability, we identify Quantitative Trait Loci (QTL) in the host plant associated with 18 phenotypic traits in the parasite as a resource for the community. Finally, we leverage the large and diverse dataset to understand fundamental features of the parasite, independent of host genotype, revealing aspects of the life cycle which are pseudo-deterministic as well as local, deleterious interactions between co-infecting parasites. Given that plant-parasitic nematodes cause an estimated $100 billion in agricultural damages per year, these insights are contextualised in a global challenge driven by plant-parasitic nematodes.

## Introduction

Host-parasite interactions are dynamic systems. Often, multiple invaders will interact with the host at a given point in infection, and this will change over the course of infection. Host biology involved in blocking the establishment (1) and spread (2, 3) of invaders is well described. However, framing parasitism from an invader-centric view reveals a different set of challenges, less well studied (4): e.g. inter-individual differences in biology, choice of infection site, outcomes of local host-parasite interactions, and cooperation/competition between conspecifics.

Plant-parasitic cyst nematodes (*Heterodera* and *Globodera* spp.) are an excellent model to address these questions because: several plants, including major food crops, host cyst nematodes and are damaged by them; multiple parasites interact with the host at the same time; and the sedentary nature of host and parasite lends themselves to longitudinal studies. Importantly, these nematodes alter plant development, while the plant alters nematode development, such that individual outcomes of parasitism vary dramatically (5). For example, after destructively migrating through host tissue, second-stage juveniles (J2) select a single cell from within the vascular cylinder which is induced to form a large, syncytial, feeding site from the fusion of hundreds of adjacent cells. Concurrent with this selection, the nematode becomes sedentary and starts non-destructive feeding from the syncytium for the remainder of its life (several weeks). The feeding site is the sole source of nutrition for the developing nematode, and the early stages of feeding are concurrent with post-embryonic sex determination of the nematode (5), highlighting both the plasticity of development and the importance of the early choice.

Here we take advantage of recent advances in deep-learning-based phenotypic analysis (reviewed in (6)) to derive biologically relevant information (i.e. traits or infection patterns) from timelapse videos of tens of thousands of individual *H. schachtii* parasites, infecting thousands of *A. thaliana* hosts, across hundreds of genotypes. By studying individual parasitic entities infecting in concert, we revealed a greater extent of host genetic control of parasite development, as well as fundamental, host-genotype-independent, features of parasite biology.

## Results

### Low-cost 3D-printed custom hardware system and Deep-learning-powered software for semi-automated trait analysis of *H. schachtii* infected *A. thaliana*

To accelerate and expand the scope of plant-nematode phenotyping from manual and heuristic-based quantification (7), we designed, 3D printed, and manufactured a novel hardware system (see Figure S1 for hardware comparison). This new hardware can illuminate, manoeuvre, and image *H. schachtii*-infected *A. thaliana* plants in 5-cm petri dishes. In brief (Figures 1A & S2), petri dishes loaded onto a receiving arm are individually selected and oriented under the imaging tower by an indexed wheel, automatically. The sequence for time-lapse imaging includes: illumination on, red-green-blue (RGB) image capture, write verification, unique time-stamp, and illumination off. The petri dish is then automatically deposited into the returning arm while the next petri dish is selected. In this way, each machine can process 1 petri dish every ∼7 seconds, at a cost of ∼£200.

**Figure 1.**
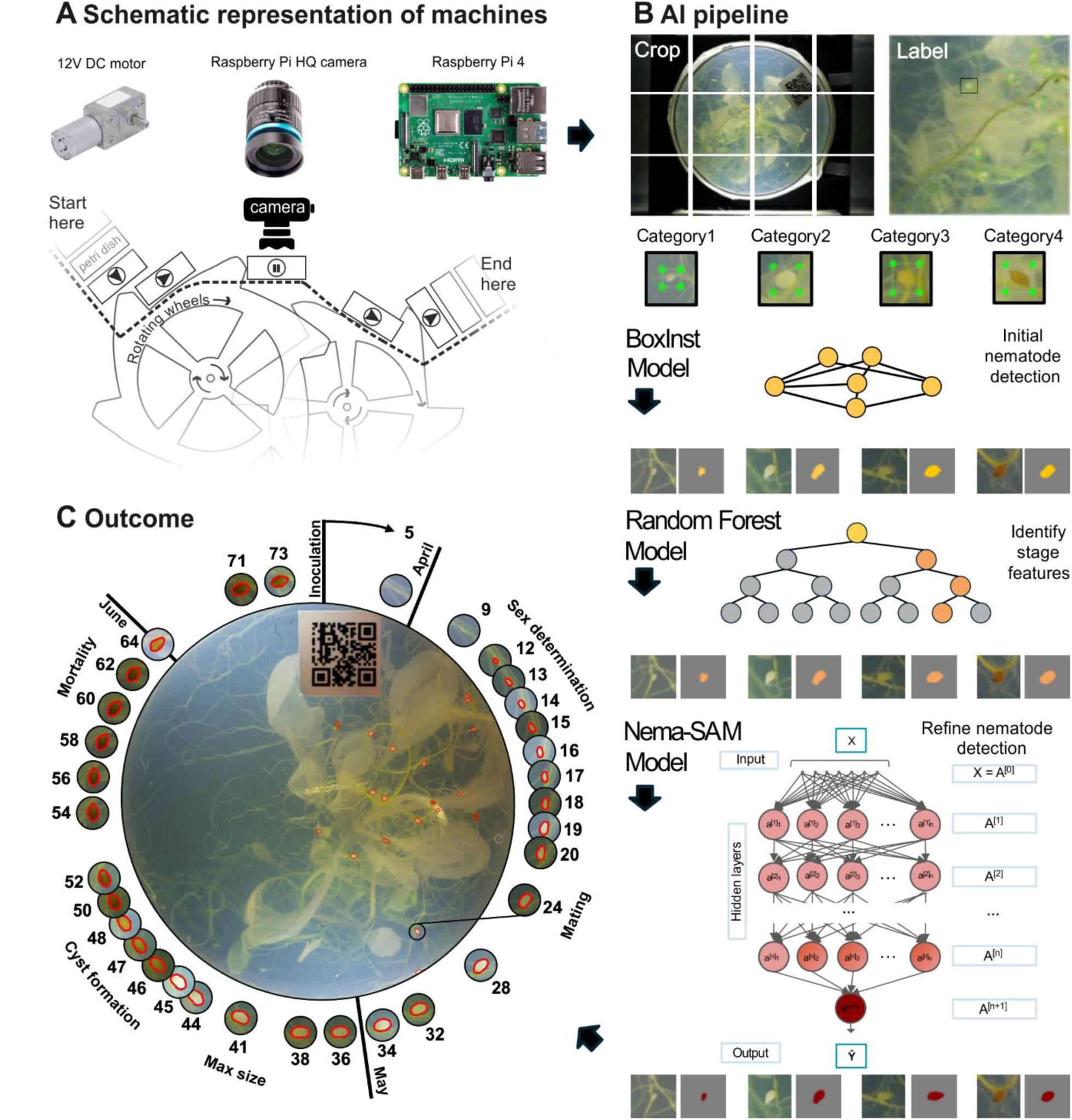
Semi-automated analysis of *H. schachtii* infected *A. thaliana* using multi-stage deep-learning-based analysis. **A**) A schematic representation of the workflow for the custom 3D printed hardware to illuminate, manoeuvre, and image 5 cm petri dishes containing *H. schachtii* infected *A. thaliana* (hardware components include 12V DC motor, Raspberry Pi HQ camera, 15 mm telephoto lens, and *Raspberry Pi* 4 computer). See Figure S2 for a more detailed workflow. **B**) Summary of the multi-stage DL-powered trait analysis pipeline, including BoxInst, Random Forest, and Nema-SAM, trained for accurately detecting nematodes from time-lapse image series: (1) the BoxInst model was used to identify nematode instance in the acquired images (categories 1-4 refer to J3/J4 females, Adult females, Adult females tanning, and Cysts respectively); (2) the Random Forest was employed for identifying morphological features such as the shape of nematode; (3) the Nema-SAM model was utilised to refine the object detection so that nematode objects were accurately delineated to enable monitoring of developmental stages of parasites over time. See Figure S5 for a more detailed workflow. **C**) An example timepoint of an infected plant where nematodes are detected (red outlines). A given nematode (highlighted with a black circle) is illustrated at every time point in cropped circles around the perimeter. Approximate life stages and days post-infection are shown.

We chose to apply this new imaging technology to the *A. thaliana* Multiparent Advanced Generation Inter-Cross (MAGIC) population (8) infected with *H. schachtii*. The MAGIC population is derived from inter-crossing 19 parental accessions which represent the genotypic and phenotypic diversity of the species (8). Time-lapse imaging of this population was performed to build a large and diverse dataset of infection phenotypes for deep-learning (DL)-powered analytics. We infected all 527 MAGIC lines with *H. schachtii* (population Bonn), each with approximately 20 biological replicates, in a randomised block design (Figure S3), totalling approximately 10,000 plants. Six machines were used in parallel to image all ∼10,000 plants, at 45 timepoints, over an 86 day period, resulting in approximately 400,000 raw RGB images. These, in essence, represent thousands of timelapse videos of infection dynamics (Video S1).

To accompany the new hardware, new software was also developed (see Figure S4 for software comparison to manual and heuristic-based quantification (7)). Selected images across infection (*n* = 2,028 petri dish images) were used to train a custom multi-stage DL-powered analysis pipeline, which automates nematode detection within the petri-dish region, segmentation of nematode objects, and nematode-level trait analysis (Figure 1B). In brief (Figures 1B & S5), after image preprocessing (e.g. colour and position calibration), every image was uniformly divided into 12 parts, and four categories of nematode morphology, which represented key parts of a parasitism cycle, were manually annotated (i.e. J3/J4 females, Adult females, Adult females tanning, Cysts). These labelled patches (including 42,659 annotated nematodes) were used to train a BoxInst model (9), which performed instance segmentation of nematode objects using a bounding-box supervision-based approach. The segmented instances (i.e. recognised nematodes) were examined by a trained Random Forest model (10), which classified identified nematodes into the four phenotypic categories based on nine predefined colour features (RGB, HSV, and LAB colour space). Finally, a customized neural network, Nema-SAM (based on FastSAM (11)), was trained to refine the delineation of nematode objects by leveraging deep semantic features, enabling more precise measurement of morphological features and thus the identification of developmental stages. Comparing manual and automated analyses (Figure S6) using representative images (*n* = 380 randomly selected nematodes) across the experiment life cycle (∼38 time points, 86 days), we obtained a highly significant correlation for the nematode number (R^2^ = 0.88, *p* < 0.0001), nematode size (R^2^ = 0.75, *p* <0.0001) and nematode roundness traits (R^2^ = 0.52, *p* < 0.0001). Given that roundness is one of the most challenging traits to predict, selected lines at the top and bottom end of the distribution were repeated, and the extremes are partially replicable (R^2^ = 0.44, Figure S7). Given the large apparent environmental contribution to roundness (R^2^ = 0.52) the between repeat correlation of R^2^ = 0.44 is to be expected.

Based on the position of each identified parasite, we connected parasitic entities between images in the timelapse videos, enabling us to monitor phenotypic changes of a given parasite during the 86-day period. Leveraging this capability across the longitudinal dataset, we quantified growth dynamics of all identified parasites over time, creating the first dynamic repository of individual pathogens, with tens of thousands of parasites infecting thousands of hosts (Figure 1C).

### Plastic development of parasitic nematodes infecting the *A. thaliana* MAGIC population

To interrogate the nature of *H. schachtii* infection of *A. thaliana*, we extracted nematode phenotypic traits from the timelapse-videos. These phenotypes are broadly categorised into the static (i.e. in an acquired image at a given time point) and dynamic (i.e. phenotypic changes over time). Example static phenotypes include, but are not limited to (Table S1 and S2), the number of nematodes (Figure 2A), their size (Figure 2B), their colour (Figure 2C), and their shape (Figure 2D). In total, we measured these four phenotypes for each of 69,054 individual parasites on 7,738 hosts at 41 days post-infection (dpi) and the same four phenotypes (as well as the ratio of 64/41 dpi number, the number of cysts, and the ratio of encysted nematodes) for 82,487 parasites on 7,962 hosts at 64 dpi. For dynamic phenotypes we first recapitulated the growth curves of individual parasites across the timelapse-videos, and then extracted dynamic phenotypes such as, but not limited to (Table S1), maximum growth rate (the slope of the tangent of the steepest point of the growth curve, Figure 2E), maximum size (the height of the curve, Figure 2F), and their time to maturity (defined as the end point of the fast growing stage, Figure 2G). In total, we measured 8 dynamic phenotypes on the highest-quality subset of the timelapse-videos (3,405 plants from 430 lines), representing over one million parasite measurements across all timelapse-images (1,237,820).

**Figure 2.**
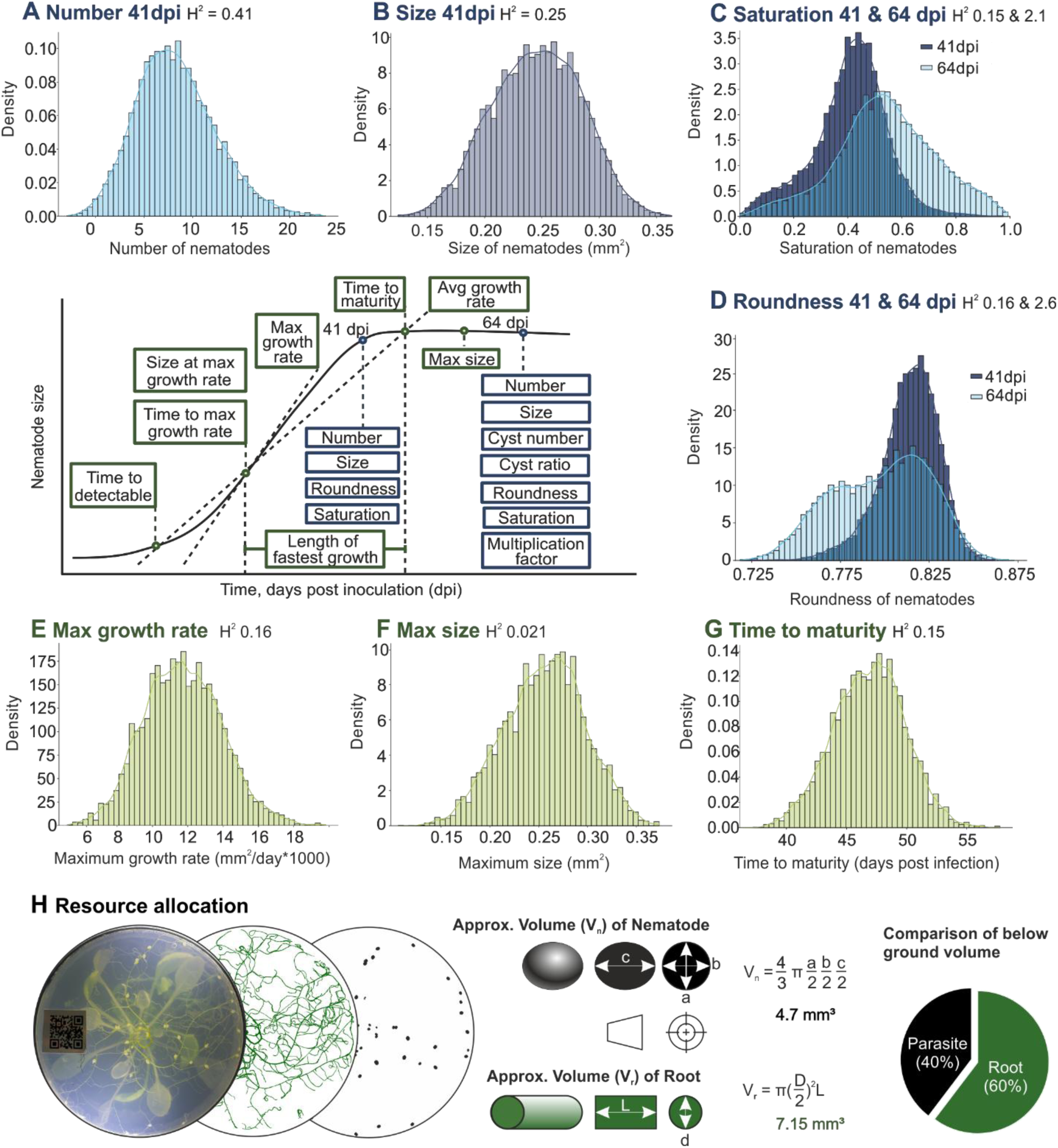
Distribution of static and dynamic nematode traits in the *A. thaliana* MAGIC population. Displayed around a central archetypal growth curve are density distribution plots (Density = Frequency/(Total data points * Bin width)) for selected static and dynamic phenotypes. **A-G**) Distributions of adjusted means (BLUEs) are shown for each selected phenotype. dpi = days post infection, H^2^ = heritability. **H** Comparison between the sum of the below-ground volume attributed to the first generation of parasitic nematode (Vn, assumes nematodes are moderately prolate spheroids where a≈b, black), to the sum of below-ground volume attributed to the roots (Vr, assumes roots are cylinders, green) at 41dpi for an example of MAGIC line 133.

These data demonstrate that the *A. thaliana* MAGIC population contains variation, typically not larger than an order of magnitude, for all measured parasite traits: including the number of successful infections as well as the development and reproduction of those individuals. Interestingly, however, heritability (i.e. plant genetic control of nematode traits) also varies. Static phenotypes generally have higher heritability than dynamic phenotypes, with the highest heritability for nematode number at 41 dpi, H^2^ = 0.408; while the lowest heritability was for maximum size, H^2^ = 0.021). Together, these data show that while dynamic phenotypes typically have larger environmental contributions than static phenotypes, variation in plant genotype has substantive contributions to variation in most measured aspects of the nematode life cycle.

The dominant majority of these nematode phenotypes are normally distributed, yet, those which are non-normal are additionally informative of the biology captured across the dataset. For example, comparing the distribution of colour saturation at 41 dpi with that at 64 dpi shows that a subset of nematodes encyst (as indicated by saturation, Figure 2C). Similarly, comparing roundness (Figure 2D) at 41 dpi with that at 64 dpi is indicative of a second generation of nematodes re-infecting the same host (see also Video S1). These distributions set upper and lower physiological limits for the species, under these conditions. Most remarkably, line133 is an outlier on both the number of nematodes infecting the plant (top 1.5 percentile) and an outlier on the size of those nematodes which result (top 4.5 percentile). We used an example from this line to estimate the physiological limit of susceptibility under these conditions, by comparing below ground volume attributed to the host with that of the parasite. Parasite volume (∼4.7 mm^3^) is of the same order as below ground volume of the host (∼7.15 mm^3^) after 41 dpi (40% and 60% respectively, Figure 2H). Given that at 0 dpi, plant root volume is estimated to be ∼4.62 mm^3^, this means that a majority (65%) of new below-ground volume post infection can be attributed to the parasite, in this exceptional case.

### Nematode development is determined by variation in plant genotype

To search for plant genetic loci which associate with nematode traits, we can take advantage of the fact that the genome sequence of each MAGIC line is already determined (8). This, therefore, enables Genome Wide Association Studies (GWAS) between nematode phenotypes and plant genotypes to identify SNPs linked with candidate genes which may contribute to the traits measured. Using this capability, we identified Quantitative Trait Loci (QTL) in the host plant associated with 18 phenotypic traits in the parasite as a resource for the community (Figure 3, S8, and Table S1), including static and dynamic traits. For example the static trait nematode number at 41 dpi (Figure 3A) is associated with several QTL on chromosomes 1, 3, 4 and 5 – with the major QTL being Chr.1: 25477452-25915205, Chr.1: 26705856-27697579, and Chr.5: 4042687-4685737. Each major QTL contains 34/131, 97/341, and 68/210 of the underlying genes variant in the population screened, potentially including causal genes. An example haplotype analysis for SNP 27642477 under the broad peak on chromosome 1 shows three genetic variants present in the population, two comparatively resistant and one comparatively susceptible (Figure 3Aiii). Interrogating the pedigree shows that, in this case, the minor allele associated with resistance (n=35 MAGIC lines) is exclusively derived from the parental line Can-0, the minor allele associated with susceptibility (n=17 MAGIC lines) is exclusively derived from Zu-0, while all other MAGIC and parental lines (including Col-0) harbour the major allele associated with resistance (n=324 MAGIC lines). An F1 hybrid between a MAGIC line harbouring the major allele associated with resistance (M373) and a MAGIC line harbouring the minor allele associated with susceptibility (M166) suggests that in this case susceptibility is dominant. In addition, we showed that GWAS over time for nematode size can be used to identify a transient effect on nematode size (Figure 3B). We identified that small differences in the distribution of nematode sizes over time (Figure 3Bi) are associated with very large differences in the strength of the association with plant genotype (Figure 3Bii). In total, each Arabidopsis chromosome harbours several discrete and adjacent QTL for a variety of static and dynamic traits nematode traits (Figure 3C).

**Figure 3.**
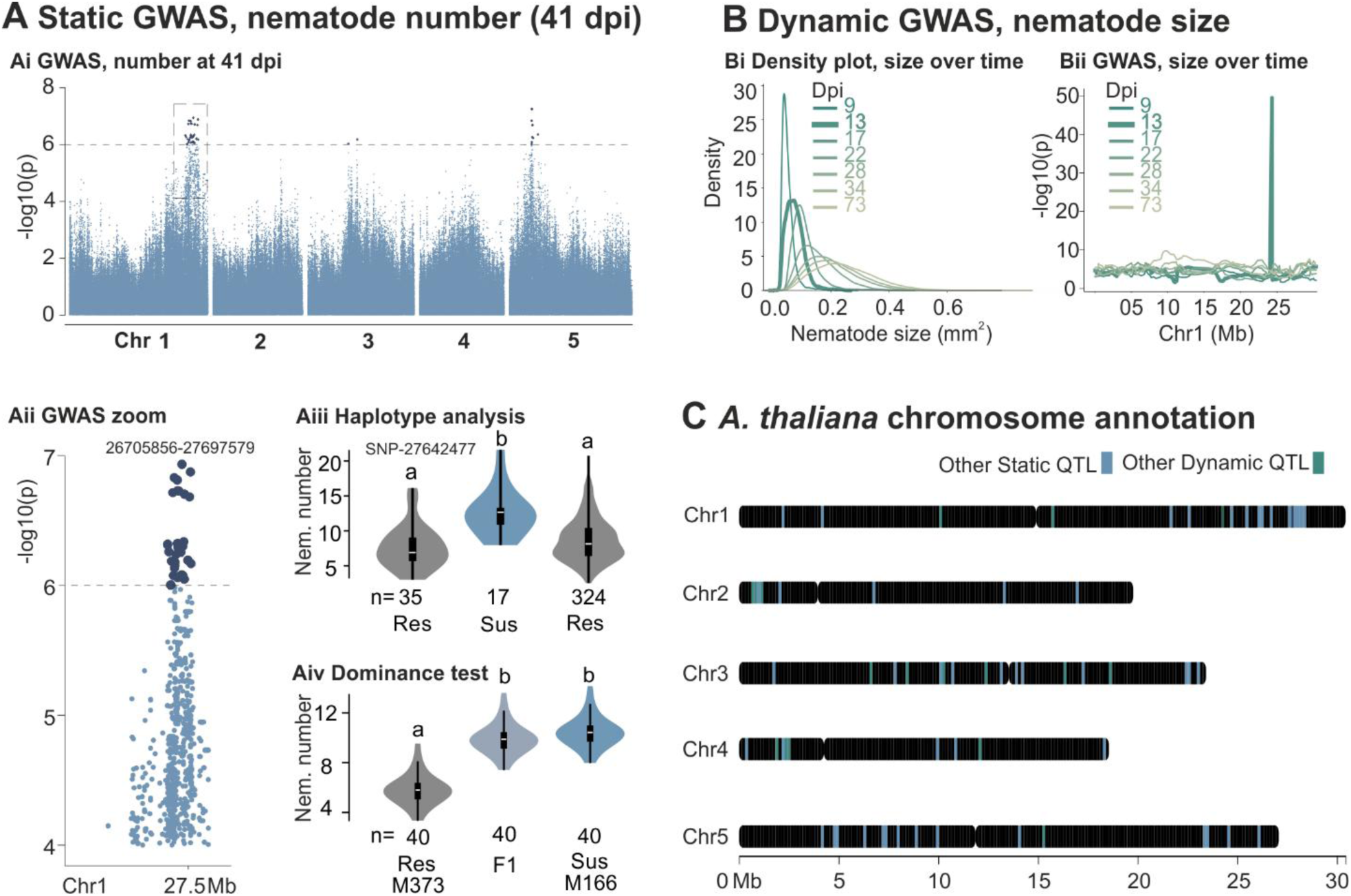
Nematode development is determined by variation in plant genotype. **A)** Ai and Aii, Genome-wide association study (GWAS) for static phenotype nematode number at 41 days post inoculation (dpi). Aiii, Haplotype analysis for SNP 27642477, three haplotypes are present, n=number of individuals in the screen harbouring a given haplotype. Lower case letters indicate significant differences at p<0.05. Aiv, result of cross between comparatively resistant (Res) parent (line M373) and comparatively susceptible (Sus) parent (line 166) results in F1 which mimic susceptible parent. n=number of replicates, lower case letters indicate significant differences at p<0.05. **B**) Bi Density plot of nematode size over time. Bii GWAS of nematode size over time. **C)** *Arabidopsis thaliana* chromosome-wide annotation based on summary of GWAS for static and dynamic phenotypes (see Figure S8 and Table S1 for details).

### The nature of *H. schachtii* infection of *A. thaliana*

Given the breadth of phenotypes analysed, for tens of thousands of nematodes, on hundreds of plant genotypes, we can determine the relatedness of traits and thereby infer the nature of *H. schachtii* infection of *A. thaliana* (Figure 4A). For example, pairwise correlations between static traits show that, across the ∼500 genotypes tested in this paper, there is a global positive association between nematode number and size (R2 ∼0.25, p<0.001, Figure 4Ai and Figure S9) - i.e. at the same inoculation density, the more nematodes that infect a plant the better those nematodes perform individually.

**Figure 4.**
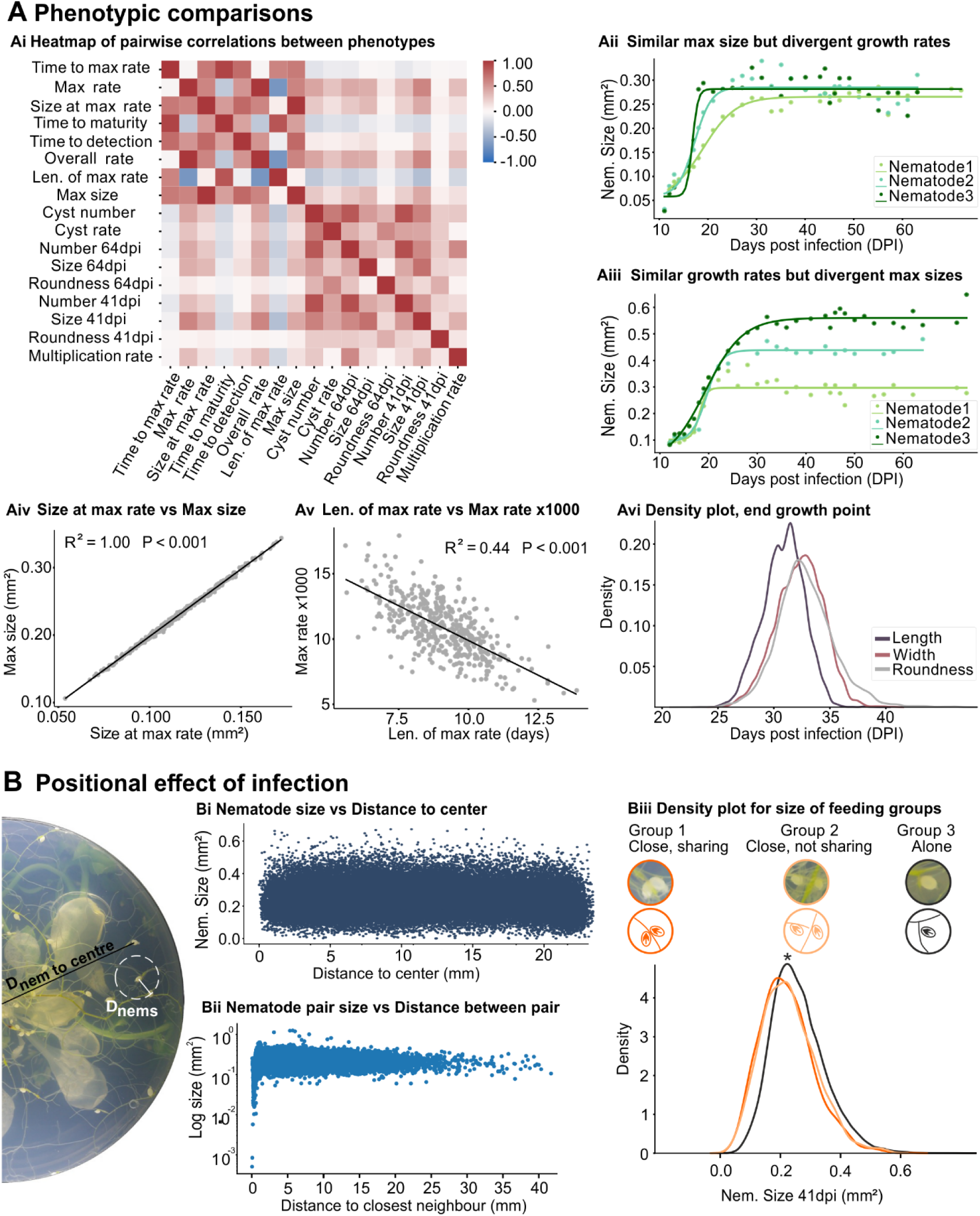
The nature of *H. schachtii* infection of *A. thaliana*. **A**) Phenotypic comparisons. Ai, pairwise correlation between each phenotype extracted from image series. Aii and Aiii, example growth curves showing opposing co-variance. Aiv, correlation between Max size and size at maximum growth rate. Av, Correlation between length of maximum growth rate period and maximum rate. Avi, Density distribution plot (Density = Frequency/(Total data points * Bin width) of point of last change in Length (blue), Width (red), and Roundness (Grey). **B**) Positional effect of infection. Left, example image with illustrations to explain measurements of distance of each nematode to the centre of the plate (black, as per Bi), and the distance to the closest neighbour (white, as per Bii). Biii, Density distribution plot (Density = Frequency/(Total data points * Bin width) of nematode size for three groups: Group 1 (nematodes that are close (<0.915 mm distance between the two), and share a root), Group 2 (nematodes that are close (<0.915 mm distance between the two), but do not share a root), and Group 3 (nematodes that are alone (nearest nematode >0.915 mm)). Asterisk indicates significant difference between Group 3 (*n* = 60,933) and Groups 1 (n = 2,540) and 2 (n = 1,031) ANOVA and Tukey HSD, and pairwise T-test, (*p*<0.001).

Interestingly, and even though individual parasites exist with similar maximum size but very different growth rates (Figure 4Aii), and others exist with dissimilar maximum size but very similar growth rates (Figure 4Aiii), there is a negative correlation between max growth rate and the duration of the fastest growth period (Figure 4Av, R2 = 0.44). i.e. those nematodes which grow faster, stop growing sooner. Indeed, for a well-fit sigmoid curve, the size at maximum growth rate is necessarily correlated with maximum size (R2 = 1.0, Figure 4Aiv). Taken together, these data suggest that maximum size has deterministic elements. Interestingly, how these nematodes put on this size appears to be broadly allometric: the point at which growing nematodes stop gaining length (mean approximately 30 dpi) is before they stop gaining width (mean approximately 34 dpi), and thereby roundness (Figure 4Avi).

By leveraging the positional information of each parasite on each host, we can also interrogate both the interaction between parasite and host, as well as the interactions between parasites on the same host. First, we sought to determine whether, at the organismal level on each host, there are parts of the root system which enable greater nematode nutrition. Given that seeds were sown approximately central to each petri dish, the distance of each nematode to the centre of the petri dish is therefore a rough proxy for position on the tap root system (Figure 4B). Remarkably, distal parts of the root system produce the same sized nematodes as those most proximal to the aerial parts of the plant (R^2^ = 0, Figure 4Bi).

Locally, however, there is an effect of parasite position. Plotting pairwise distance between any two nematodes on the same plant reveals a negative skew at approximate distances <50 pixels (0.915 mm, Figure 4Bii). To determine the underlying nature of the skew, we manually annotated 64,504 nematodes into one of three groups: Group1, nematodes in close proximity and infecting the same root (2540); Group2, nematodes in close proximity but are infecting different roots (1031); and Group3, no nematode in near vicinity (60,933 number, >50 pixels (0.915 mm)). Lone nematodes are significantly larger, across the whole distribution, than those close to other nematodes - whether they share a root or not (Figure 4Biii, ANOVA and Tukey HSD, pairwise T-test, P<0.0001). Interestingly, this local negative effect of proximity is in the context of a global positive association between nematode number and size (Figure 4A and Figure S9). These appear to be independent phenomena because stratifying the approx. 60,000 nematodes in Figure 4Biii by the number per plant – into those in the top half of global infection density, and those in the bottom half – reproduces an effect of equal magnitude in each half (Figure S9).

## Discussion

### Technological advances in the scale and scope of phenotyping

Bespoke imaging platforms like that developed herein, when coupled with DL-powered phenotypic analysis pipelines, can have a transformational impact on discovery by increasing both scale and scope of investigation. To put the scale into perspective, four experimenters operating six imaging machines in parallel provides an approximate theoretical imaging speed of 1 plant per second - in our hands, 10,000 plants were routinely processed in under 4 hours. Manually detecting nematodes on 10,000 plants may take the same experimenters over 80 hours (assuming ∼2 minutes per plant, without any breaks). As big as this difference seems, it is a false equivalency because the DL-powered software extracts multiple traits, many of which are unmeasurable by humans at scale (e.g. static size, shape, and colour measurements, or phenotypic changes of any kind over time). As such, this scale enables holistic and dynamic pathology: in this context, holistic refers to the measurement of all individual parasitic entities infecting the plant, while dynamic refers to a longitudinal analysis of said individuals over time.

Some traits are uniquely able to describe aspects of the life cycle (e.g. growth, maturation, encystment), and yet they have remained unstudied because they were, to all intents and purposes, manually unmeasurable previously. With our segmentation-approach being largely agnostic to these differences - for example, size, shape, and colour are inherent in the nematode-level detection (number) - these historically most challenging traits may now hold the greatest potential for novel biological discoveries. Some traits remain challenging. For example, roundness is both the most difficult to predict and most difficult for us to interpret. It is not clear why or how nematode roundness (independent of nematode size) has components under plant genetic control. It seems that the phenotypes described in this paper are distinct from sex-related differences because comparison to Anwer et al., (13), who performed a GWAS of *A. thaliana* variation on sex ratio of *H. schachtii*, shows no overlap. Taken together, the implication is therefore that, in addition to expanding scale, this approach expands the scope of research into a group of pathogens which are threatening global food security.

Due to the rapid progress of AI-related techniques, the AI-powered solution presented here will inevitably evolve in many areas in future developments. For example: 1) enhanced imaging diagnostics of nematodes, which can be enabled by combining spectral analysis and multi-granularity hierarchical attention mechanisms (14) with vision transformers (15), distinguishing nematode growth stages and species from background signals (e.g. roots and soils), with an increased accuracy; and 2) digital twins for simulating nematode-plant interactions and infection dynamics under varying conditions (16), unlocking additional orders of magnitude increases in scale.

### What does it mean to be resistant to plant-parasitic nematodes?

Given that: i) upon completing infection, cyst nematodes can remain dormant in the soil for decades, and ii) that plants exhibit a parasite tolerance threshold, above which damage is manifest (18, 19), functional resistance is that which reduces nematode reproduction factor (the initial population of nematodes in the soil after harvest divided by the population of nematodes in the soil before planting). While it has been known for decades that the reproduction factor is multivariate (20), many of these traits rarely feature in the discussion on resistance, and there has been little progress in understanding some traits which both vary dramatically between plant accessions and which contribute to reproduction factor: nematode size, growth rate, generation time, etc. In particular for parasites like *H. schachtii*, which are capable of multiple (up to 5) generations on their host in a single season (21), genes which control the rate of growth and reproduction can have a greater impact than genes which control the number of nematodes, because generations are multiplicative. So while classical immunity can be important for cyst nematodes (22), clearly the diversity of genes that drives these phenotypes is likely greater than those involved in immunity alone, and indeed the evidence is clear that most resistance genes against Heterodera spp. are non-NLRs (23, 24). Therefore, the provided expansion in scope may also be necessary to understand the full gamut of what it means to be functionally resistant to plant-parasitic nematodes in an agricultural context (22).

### Future exploration of host genetic control of parasite traits

Hardware and software have been made open source, all images are available in an open repository, all derived traits are available in Table S2, and using this information the *A. thaliana* genome is now annotated for QTL across 18 traits. Open access to these data at each stage of generation and analysis represents valuable resources for future research which can be built upon with new detection algorithms, new analyses, or deeper exploration to identify host genetic control of parasite traits. The limited overlap between QTL, taken together with the observation that static phenotypes typically correlate better with other static phenotypes than they do with dynamic, and vice versa, highlights the additional value a longitudinal analysis of parasitism holds for discovery. Parallel work on this same data set has shown that in the dominant majority of cases the quantity of root before infection provides no rational prediction for the number of nematodes post infection (i.e. there are as many lines where root area positively correlates with nematode number as there are lines were it negatively correlates with nematode number Zhou et al., in press). Nevertheless, future work would likely integrate these datasets for a holistic view of covariance between plant and nematode traits to refine/triage QTL predictions.

### Parasite control of host defences and resource allocation

While DL-powered phenotyping of pathologies has contributed to our understanding of the genetic basis of resistance in general (e.g. Figure 3, and reviewed in (29)), its contribution to our understanding of pathogen biology is, to the best of our knowledge, absent. Here we leverage the dataset generated to also understand fundamental features of the parasite.

For example, in spite of the diversity of host genotypes tested, derived from MAGIC parents chosen to represent the genotypic and phenotypic diversity of the species (8), we found no single line which is entirely resistant to nematode infection. This highlights the profound ability of nematodes to manipulate and/or avoid host defences, and is consistent with the literature. What is less well studied is whether and how nematodes manipulate host resource allocation (30). At the other end of the spectrum, in the limit of susceptibility for this population, we found that the volume of 1st generation nematodes below ground can be similar to root volume below ground at 41 days post-infection. Given that in this experiment, plants had 26 days to accumulate volume below ground before nematodes were inoculated, this is a remarkable ability to dominantly redirect nutrient allocation below ground. Evidently, this observation was made under replete nutrient conditions, and in a binary interaction. So while it is unlikely such staggering manipulation will manifest in more natural agroecosystems, this unexpected finding nevertheless highlights the substantive extent to which parasitic nematodes can rewire plant resource allocation below ground. While it remains to be determined whether, in this genetic background, nutrients are also re-directed above ground, it seems unlikely given that leaf area has also been used as a robust proxy for (in)tolerance to nematode infection across a wide diversity of *A. thaliana* genotypes (18). Combining root, leaf, and nematode measurements in one study would be required to fully describe the extent to which the nematode can alter plant nutrient resource allocation in general.

In analysing a diversity of lines, we also identified host genotype-independent features of parasitism. The first example is conceptualised as deterministic plastic development. i.e., that while there is tremendous plasticity in the growth and development of individual nematodes, some aspects are pseudo-deterministic. For example, we have shown that parasites which have higher maximum growth rates (“live fast”) tend to have a shorter duration of their fastest growth period (“die young”). This is reminiscent of the so called “rate of living theory” (31). So while there are exceptions to the rule (e.g. Figure 4Aiii), some 44% of the variance in duration of the fastest growth period is described by the steepness of said period. Therefore, we conclude that once a successful feeding site is established and nematodes are engaged in peak nutrient consumption from the host - they often maintain an established trajectory to a maximum size that is in large part (44%) determined by that rate of growth. Interventions which alter or disrupt these earliest stages of nematode growth and development, as with those that block feeding site establishment altogether (22), will also likely have an outsized impact on reproduction factor.

### Short-distance deleterious “communication” between parasites

Finally, we show that where these parasites infect the host impacts their growth. We observed a local effect of position: nematodes near one another tend to be smaller than nematodes without a near neighbour. This effect is, counter intuitively (32), in the context of a global positive association between nematode number and size, but independent thereof (Figure S9). Given that feeding sites of adjacent nematodes can fuse to a single contiguous syncytium (33), an intuitive interpretation is competition. However, this seems unlikely given that: i) the entire distribution shifts - there is no winner; and ii) unexpectedly, even those nematodes which are close to one another but not sharing the same root still suffer from the effect. Therefore, the most parsimonious explanation is a diffusible signal. These data require some degree of “communication” - in the broadest sense - between parasites and/or feeding sites, outwith the root network. Options may include a nematode-derived signal (e.g. some sort of pheromone), a plant-derived signal (e.g. some sort of damage signal), or some combination thereof. Future work leveraging parallel root detections could determine the connectedness of the different roots infected by proximal nematodes to determine whether this enhances the observed effect. This would require additional DL-powered models to detected and define the details of root architecture in this system, under nematode burden.

In summary, DL-powered phenotypic analysis enabled us to extract meaningful biological information from timelapse videos of tens of thousands of individual parasites, infecting thousands of hosts, across hundreds of genotypes. This approach allowed us to understand a greater extent of plant genetic control of parasite traits, describe parasite control of host resource allocation below ground, as well as identify fundamental (plant-genotype-independent) features of parasite biology. Digitising the infection process in this way, and making the timelapse-videos open source, will allow others to expand upon the resources as new analysis pipelines are developed.

## Methods

### Phenotyping platform

#### Hardware and control code

The bespoke hardware imaging machine was designed either via Onshape (PTC Inc.), or Autodesk 123D design, and exported in STL format. STL files were sliced in CURA (Ultimaker) to generate G-code instructions for the Ultimaker S3 printer, fitted with one AA, and one BB, 0.4 mm printing cores (Ultimaker). All the components of the imaging machine were printed utilising Tough Polylactic Acid (Ultimaker) with the following printing settings: infill density 40% (triangular pattern), layer height 0.2 mm, wall thickness 0.8 mm. A full hardware parts list can be found on: https://www.hackster.io/nemaspotters/low-cost-high-throughput-imaging-of-universal-petri-dishes-b5d6e9.

Machine operation was controlled via a Raspberry Pi 4B running Raspberry Pi OS 11. A 12 MP HQ camera (Raspberry Pi) was fitted with a 16 mm telephoto lens (Raspberry Pi). Illumination was provided by two opposing 12 V LED Chips (Zerodis). Both destacking and restacking disks were rotated via a 6 RPM 12 V DC geared motor (Lazmin) connected to a 130MXL timing belt (Contitech), which joined the rotation of the two wheels. A hall-effect sensor (Waveshare) paired with permanent magnets (Deryun) was integrated into an indexing wheel to position petri dishes precisely under the camera. The operation of the imaging hardware was controlled using custom Python scripts (https://github.com/Carinawei97/MAGIC-screen-to-cyst-nematodes.git).

Machine operation was controlled using Python (version 3.11.2). The motor was activated by default at startup. Hall effect sensor readings were stored in a list, and detection of a magnet in three consecutive reads resulted in deactivation of the motor via relay 4 and initiation of the imaging sequence. Light was turned on through relay 3. Image acquisition was performed using the libcamera-still (version 0.3.0) command, while the Python script monitored for completion of capture. Captured images were renamed with a Unix timestamp and stored in BMP format. Following acquisition, illumination was deactivated and the motor reactivated. This cycle was repeated until the process was manually terminated by the user. The full code can be found at: https://github.com/Carinawei97/MAGIC-screen-to-cyst-nematodes.git.

#### AI code

The AI workflow was developed using Python 3.8.19 and executed on a Microsoft Windows 10 workstation featuring an NVIDIA RTX 3060 GPU (12 GB memory) and an Intel Core i7-10700K CPU (32 GB memory). During the preprocessing stage, the *WeChatQRCodeDetector (*https://github.com/opencv/opencv_contrib/tree/) function from the OpenCV library was utilised to detect and decode embedded QR codes, facilitating automated image indexing and metadata retrieval. For image calibration and alignment, a scale-invariant feature transform (SIFT) (34) algorithm was employed using the *SIFT_create* function in OpenCV, which provided robust feature matching across varying imaging conditions. For nematode developmental stage classification, a Random Forest model was developed using the *RandomForestClassifier* module from the Scikit-Learn library (35). The model incorporated phenotypic descriptors to differentiate between various nematode life stages. Deep learning-based detection and segmentation were carried out using the PyTorch framework (36), with BoxInst employed for bounding box-level nematode detection and Nema-SAM used for instance mask refinement. The model output consisted of accurate nematode-level detection, with bounding boxes initially generated using BoxInst and subsequently refined into high-resolution instance masks through Nema-SAM. These instance masks facilitated the extraction of individual phenotypic traits, such as morphological characteristics (including size, length, width, and roundness) and colour features (such as saturation). All trait extraction procedures were carried out using the scikit-image library (37). The full code can be found at: https://github.com/Carinawei97/MAGIC-screen-to-cyst-nematodes/tree/main/Nematode%20traits%20analysis

### Biological material and growth conditions

This study screened 19 Arabidopsis natural accessions (MAGIC parents) and 527 MAGIC recombinant inbred lines (RILs), with seeds provided by the Nottingham Arabidopsis Stock Centre (NASC). Transgenic lines and crosses developed during this project were also used to additionally validate findings. T-DNA lines were also obtained from NASC. All phenotyping experiments were conducted using the imaging platform from this study.

Arabidopsis seeds were surface sterilised using a 20% (v/v) household bleach solution (Parazone) or chlorine gas. In the first method, the seeds were rotated for 20 minutes in the 20% (v/v) bleach solution with a Grant PTR-35 multifunction tube rotator (Grant-Bio). The bleach was removed, and the seeds were washed seven times with autoclaved de-ionised water (diH2O). For the chlorine gas sterilisation method, the seeds were placed in a desiccator with a flask containing 100 mL of sodium hypochlorite. Chlorine gas was generated by pipetting 3 mL of concentrated hydrochloric acid into the sodium hypochlorite. The desiccator was then sealed with parafilm immediately. After three hours, the chlorine gas was safely released in the fume hood, and the seed tubes sealed. Sterile seeds were covered with foil and stored at 4 °C until needed. Sterile seeds were sown on sterile standard KNOP medium containing 0.8% (w/v) Daishin agar (Duchefa Biochemie) and Gamborg Vitamin B5 solution (Duchefa Biochemie) in 5 cm deep-dish tissue culture petri dishes. Petri dishes in the large screen were placed in black trays (40 × 58 cm, Garden tray) on benches in a glasshouse, in two layers, in a randomized block design. Growth conditions were a 16-hour day (with 80 to 180 μmol m^-2^ s^-1^ light intensity) at 21 °C to 26 °C and an 8-hour night at 20 °C (Figure S2). Petri dishes for follow up experiments were grown in an incubator (MLR-352-PE, Panasonic) with the same length of lighting period, 60 μmol m^-2^ s^-1^ light intensity, at 21 °C, also with a randomized block design.

### Infection assays

*Heterodera schachtii* cysts were harvested from sterilised *Sinapis alba* plants grown on 15 cm Petri dishes containing KNOP medium supplemented with Gamborg Vitamin B5. The collected cysts were soaked in 3mM Zinc Chloride (Sigma-Aldrich) to induce hatching. Hatching was performed in a specialised 250 mL hatching jar (Jane Maddern Cosmetic) equipped with two 2.5 cm diameter plastic rings (alt-intech Tube Perspex) holding a 20 μm mesh (Sigma-Aldrich). The jars were stored at 21 °C, and hatched second-stage J2s that passed through the mesh were collected by pipetting after five days. The number of hatched J2 nematodes was estimated using a binocular microscope, and sterile Nemawash solution (0.01% v/v Tween, Sigma-Aldrich) was added to the collected suspension. The density of J2 nematodes was adjusted to 1 nematode/μL in sterile Nemawash solution before inoculating plants. After 21–26 days of plant growth, roots were inoculated with approximately 80 J2 nematodes. This was achieved by pipetting 80 μL of a well-mixed suspension (containing J2 nematodes, residual Zinc Chloride, and Nemawash solution) onto the petri dishes.

### Arabidopsis crosses

Plants were grown in controlled-environment growth chambers (16-hour day at 21 °C and an 8-hour night at 20 °C with 50 μmol m^-2^ s^-1^ light intensity) in F2 type compost (Levington Advance). Crosses were performed with 8 replicates per paternal line and 20 replicates per maternal line, pollinating 5–15 flowers per plant. Inflorescences were trimmed to synchronise flowering. Maternal buds were emasculated using fine-tipped forceps under a magnifier lamp (RS PRO), bagged to prevent contamination, and subsequently pollinated with paternal anthers. Mature siliques were harvested 14–21 days post-pollination, air-dried for 7 days, and stored at 4 °C.

## Statistical analyses

Statistical analyses were conducted in R and Python. Post-hoc comparisons between genotypes/treatments were performed with the emmeans package using Tukey’s HSD adjustment (α = 0.05). Broad-sense heritability (H^2^) was estimated to quantify the proportion of phenotypic variance attributable to genetic factors using the Distribution R package (Richard Mott, Unpublished). Variance components were extracted using the lmerTest R package. The density in the distribution plot was calculated by: Density = Frequency/(Total data points * Bin width), and the trend curve in the distribution plot represents the Kernel Density Estimate. The plot and calculation for the distribution of phenotypes were performed using Python. Visualisations included correlation heatmaps (Python, matplotlib), violin plots (seaborn), and trait distributions. Spatial distribution of nematodes was analysed using a Python script (as seen in the associated Github repository).

### Genome-wide association study (GWAS)

Phenotypic values were estimated as best linear unbiased estimates (BLUEs) using R. GWAS was performed with the MAGIC association mapping platform (8), which implements genotype reconstruction and genome-wide scans using linear mixed models (LMMs), and genome-wide significance thresholds were determined by 1,000 permutations. Complementary GWAS were conducted in R using the atMAGIC and qtl2 packages with the same imputed genotypes, enabling the incorporation of additional covariates. Manhattan and QQ plots were generated with CMplot.

### Volume estimations

Root volume was estimated based on binary masks obtained from the segmentation of Arabidopsis roots. The root system was separated into three categories: the primary root, first-order lateral roots, and the second-order lateral roots. Skeletonization of the binary masks provided the medial axis of each category, from which the total root length (*l*) was calculated. Root width was determined by measuring the shortest diameter (*d*) passing through each skeleton point, defined as the distance between the skeleton centre and the two intersection points with the corresponding binary mask boundary. The mean width of each category was subsequently used for volume calculation under a cylindrical approximation. Assuming cylindrical geometry, the overall root volume (*V*) was approximated as:

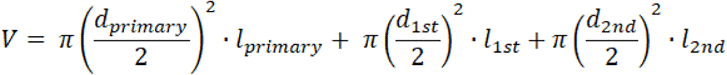

where *d_primary_*, *d_1st_*, *d_2nd_* represent the mean widths of the primary, first-order lateral, and second-order lateral roots, respectively, and *l_primary_*, *l_1st_*, *l_2nd_* denote their corresponding skeleton lengths.

The volume of nematodes was approximated using a prolate spheroid model (lemon-shaped ellipsoid). Following image segmentation, binary masks of nematodes were extracted, and morphological parameters were derived from the fitted ellipse of each mask. The major axis length was taken as the nematode body length (*c*), while the minor axis length provided the transverse body diameter (*a*). Under the assumption of axial symmetry (*a* ≈ *b*), the nematode volume (*V*) was calculated as:

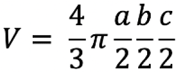

where *a*, *b* denotes the semi-minor axis of the nematode, and *c* the semi-major axis of the nematode.

### Dynamic phenotype extraction and curve fitting

Dynamic phenotypes were extracted from AI-derived nematode size, roundness, length, width, and positional data recorded across ∼50 imaging time points. To enable longitudinal tracking of individual nematodes, we developed a pairwise matching algorithm that links segmented objects across consecutive time points. First, binary masks of nematodes were generated using connected component labelling (ndi.label) and morphological refinement (e.g., border clearing). For each detected region, bounding boxes were expanded by a fixed margin to create candidate search windows. A benchmark mask from the initial frame was used as a reference, and subsequent frames were scanned for overlaps with the reference bounding box. Pairwise intersection areas were computed using the ‘get_areà function, and intersection-over-union (IoU) was applied as the matching criterion. If the IoU exceeded a defined threshold, the region was retained as a candidate trajectory. Matched regions were then relabelled to extract morphological features, including centroid position, area, perimeter, major and minor axis lengths, aspect ratio, compactness, and roundness. This allowed continuous updating of nematode trajectories over time, with trait values recorded at each time point. In cases where overlaps were ambiguous, unmatched detections were discarded to reduce false linkages. This pairwise tracking ensured that nematodes could be reliably followed across time series, providing the basis for downstream data triage, curve fitting, and growth curve analysis.

Data filtering, before curve estimation, was performed as follows: trajectories with more than five consecutive missing time points or >= 30% overall missing data were discarded (i.e. days with the no measurement for a given nematode), and a rolling filter (if a point was lower than the previous, then it was discarded if it was higher or lower than 1.5 * average ±1 dpi) was applied to remove outliers that were biologically implausible. Sigmoid growth curves were fitted to nematode measurements across time. Data on nematodes was loaded into a Pandas (version 2.3.2, (38)) dataframe. To aid sigmoidal curve fitting, three starting reference points were added for the inoculation day, equal to the area of a *H. schachtii* J2 (0.0081 mm^2^). Sigmoid curves were fitted using the curve fit function of Scipy (version 1.16.1, (39)) using the formula:

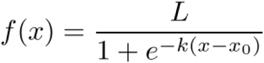

Where *L* is the curve upper bound, *k* is the growth rate, *x0* is the midpoint (½ *L*). Both scripts for data filtering and sigmoid fitting can be found on: https://github.com/Carinawei97/MAGIC-screen-to-cyst-nematodes

## Code and data availability

A full hardware parts list can be found on: https://www.hackster.io/nemaspotters/low-cost-high-throughput-imaging-of-universal-petri-dishes-b5d6e9.

Hardware and Software github - https://github.com/Carinawei97/MAGIC-screen-to-cyst-nematodes.git

Phenotype extraction github - https://github.com/Carinawei97/MAGIC-screen-to-cyst-nematodes/tree/main/Nematode%20traits%20analysis

Image data deposition – https://www.ebi.ac.uk/biostudies/bioimages/studies/S-BIAD2402

## Acknowledgements

The authors would like to thank Thiago Alexandre Moraes for discussion on growth curves and derivatives. Lawrence Percival-Alwyn and Mark Jones for advice on 3D printing and engineering, Tally Wright for block design, BLUEs and LD mapping, Hugo Tavares for QTL & GWAS, Evan Elison for providing the ptrans_220d vector, Denitsa Hristova and Natasha Yelina for advice on crossing *A. thaliana*, and Richard Mott for advice on the MAGIC population. Work on Plant-parasitic nematodes at the University of Cambridge is supported by DEFRA (125034/359149/3), BBSRC grants BB/R011311/1, BB/S006397/1, BB/X006352/1, and BB/Y513246/1, a Leverhulme grant RPG-2023-001, a UKRI Frontier Research Grant EP/X024008/1, and the Gatsby charitable foundation. Ji Zhou is supported by BBSRC AI in Plant Research Grant (BB/Y513969/1); an overseas foreign specialist project from the Ministry of Science and Technology of the People’s Republic of China: G2021145005L. Jie Zhou is supported by the China Scholarship Council program (Project ID: 202406850082). The authors would like to thank the BioMaker foundation for financial support.

## Author contributions

● Designed research

○ SW, OPK, US, JiZ, SEVDA
● Performed research

○ SW, JZ, OPK, GS, ZH, BS, CP, US, ADTB, RH, VHMdS, VCTH, TW, AGJS, AD, KvM, TB, LD, JiZ, SEVDA
● Analyzed data

○ SW, JZ, OPK, GS, ZH, TW, JiZ, SEVDA
● Wrote the paper

○ SW, JiZ, SEVDA

## Supplemental media

**Video S1. Interpolated time-lapse video for illustrative purposes.** Image series were stitched together into a video, and in between frames were interpolated using Flowframes (version 1.36.0, https://github.com/n00mkrad/flowframes) using the following settings: 10x frame multiplier, RIFE3.9 interpolation algorithm. This resulted in a video of 20 fps.

---

**Figure S1.**
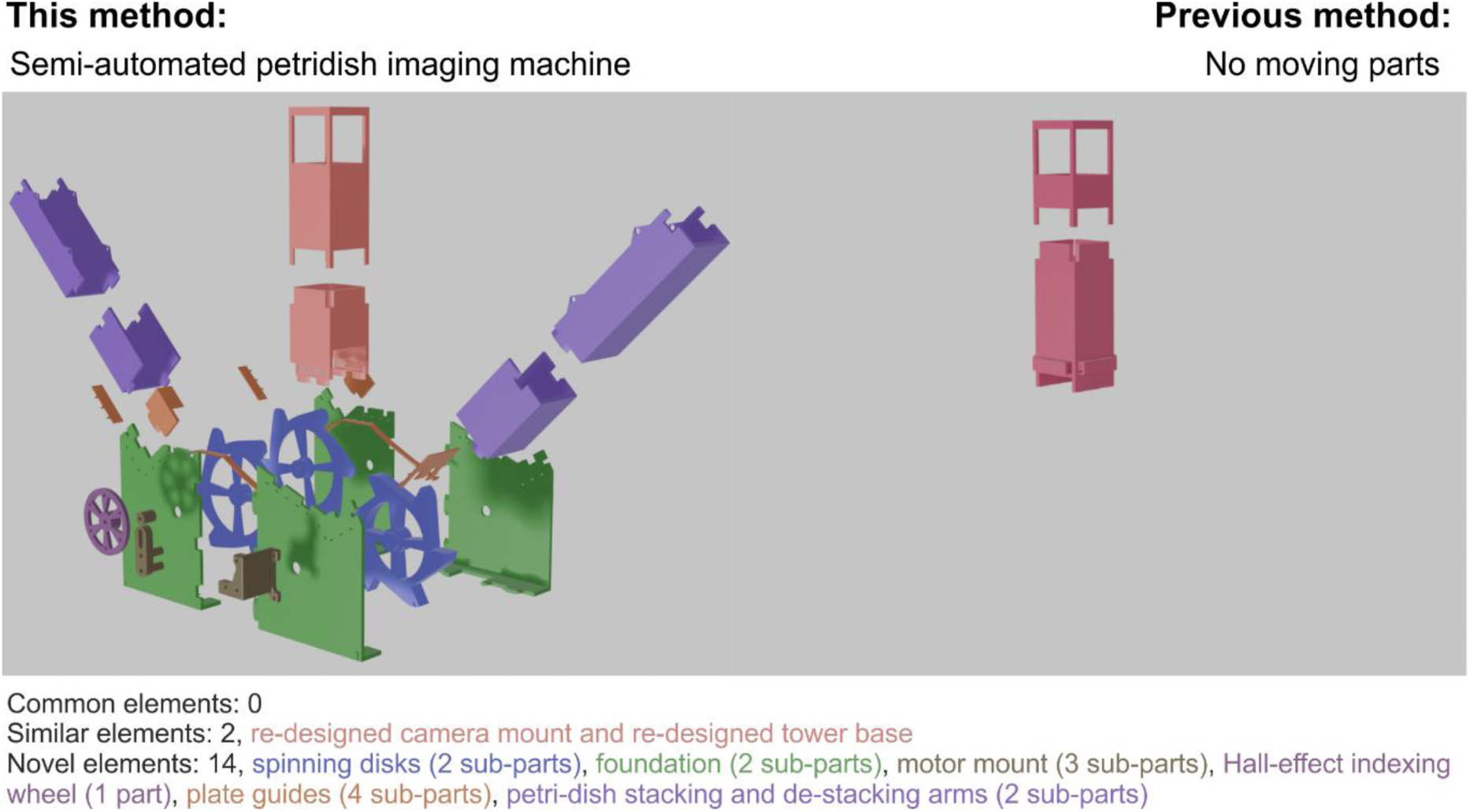
Hardware comparison between previous method (right) and this method (left). Parts are colour coded by function. There are no common 3d printed parts, two similar parts that were re-designed, and 14 new parts that were designed from scratch. The most striking difference is that the novel method is semi-automated, with several moving parts when compared to the previous method (Kranse et al,. 2022).

**Figure S2:**
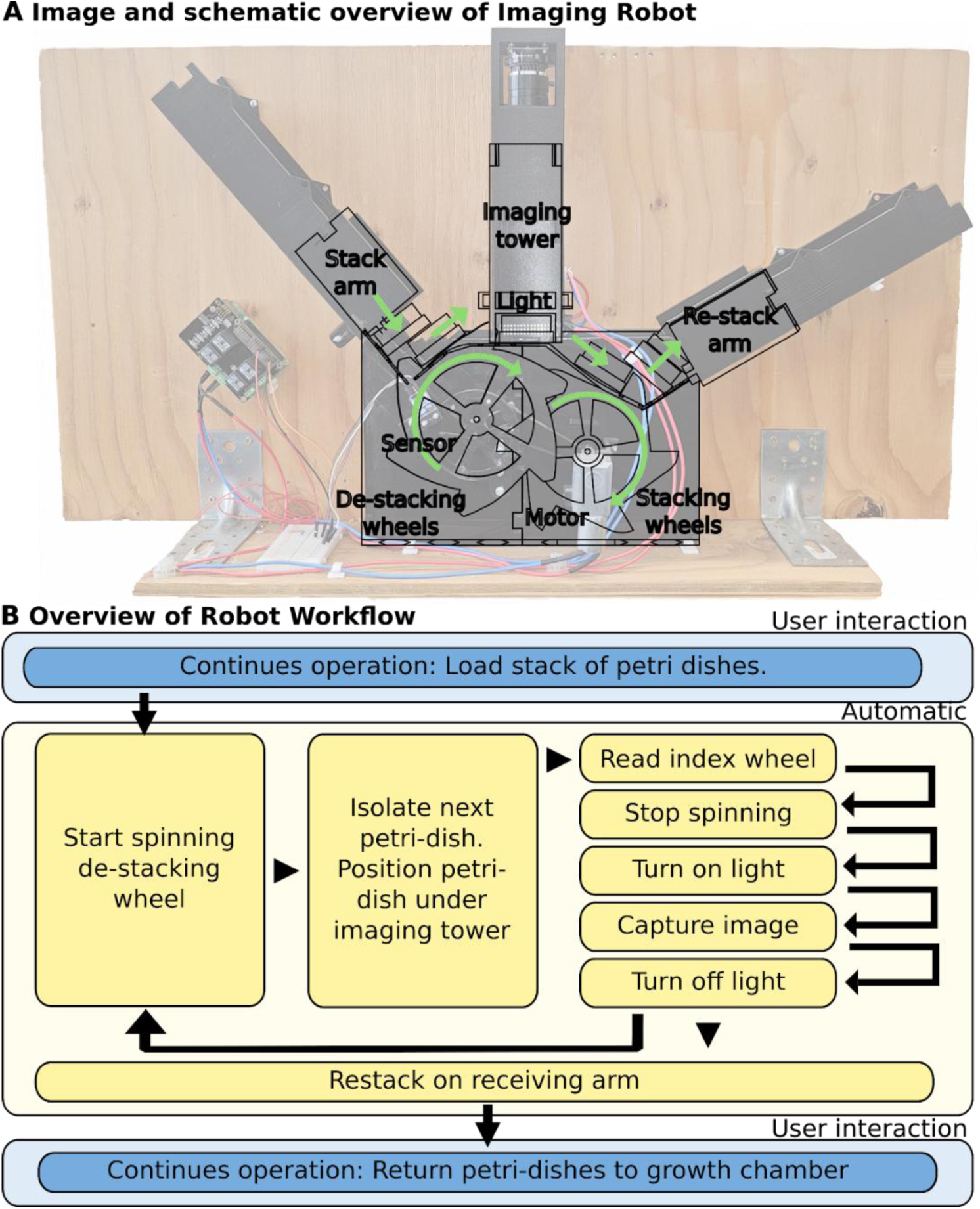
Custom 3D printed imaging hardware and workflow. **A**) Schematic representation overlaid on a photo of imaging machine. From left to right; petri-dishes are moved from the stack arm to the imaging tower via the de-stacking wheels, and later re-stacked on the other side using the stacking wheels on the re-stack arm. **B**) Overview of workflow, highlighting user initiated, and autonomous, aspects of the semi-automated imaging pipeline. The user has two operations, loading, and removing petri-dishes. Once loaded, the software starts the motor, spinning the de-stacking and stacking wheels, this isolates a single petri-dish, and places it under the imaging tower. An indexing wheel detects the potential presence of a petri-dish in the tower, and triggers an imaging sequence. When the machine restarts, the imaged petri-dish is re-stacked on the re-stack arm.

**Figure S3.**
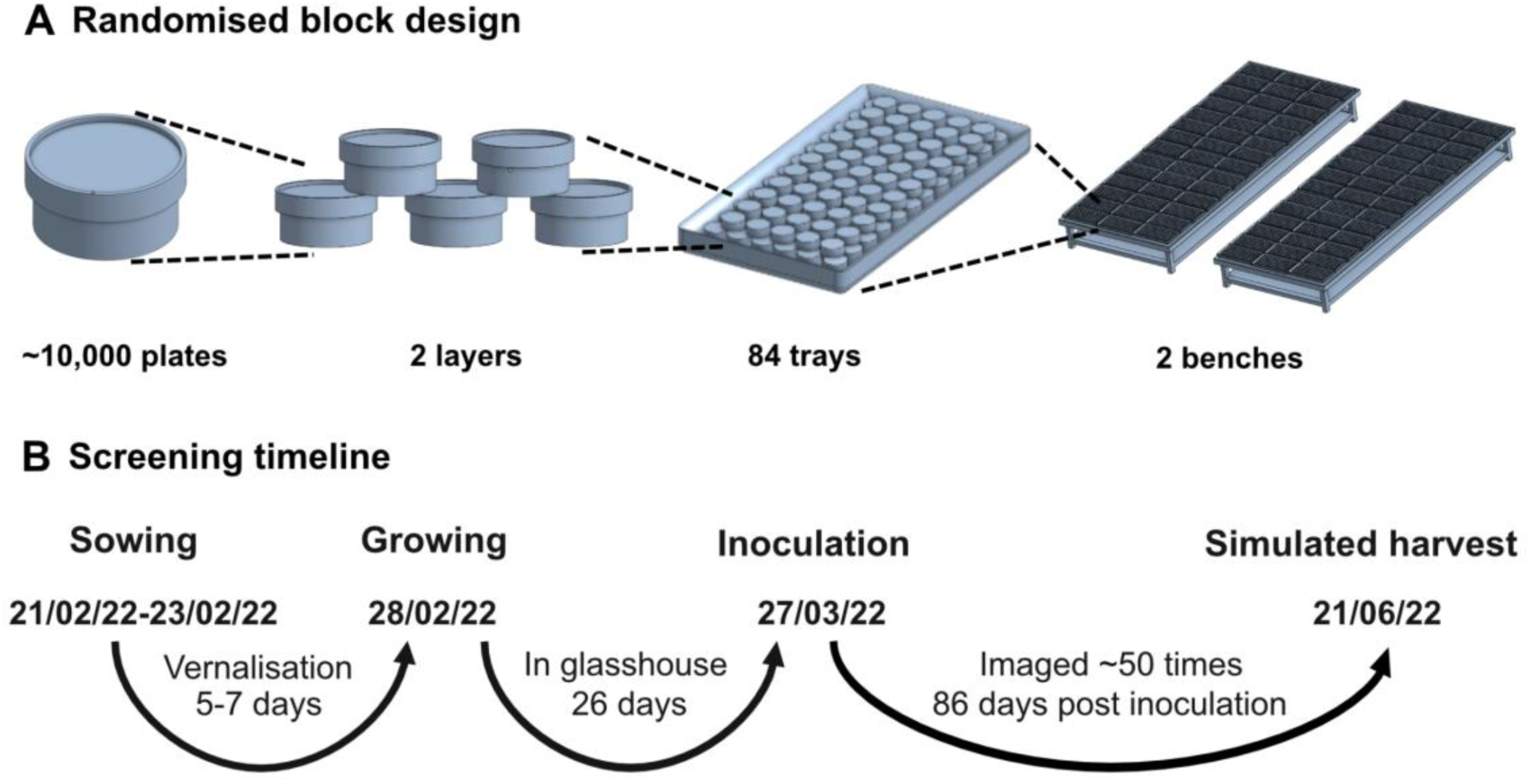
Experimental design and timeline. **A)** Schematic of the randomised block design used for the host-parasite interactions screening ∼10,000 *A.thaliana* petri dishes from 527 MAGIC lines. Petri dishes were organised in two layers across 84 trays distributed over two benches. **B)** Timeline of the host-parasite interactions screening workflow. Seeds were sown between 21/02/22 and 23/02/22 and underwent 5–7 days of vernalisation. Plants were then transferred to the glasshouse on 28/02/22 and grown for 26 days prior to inoculation on 27/03/22. Imaging was conducted approximately 50 times over the course of 86 days post-inoculation, culminating in a simulated harvest on 21/06/22.

**Figure S4.**
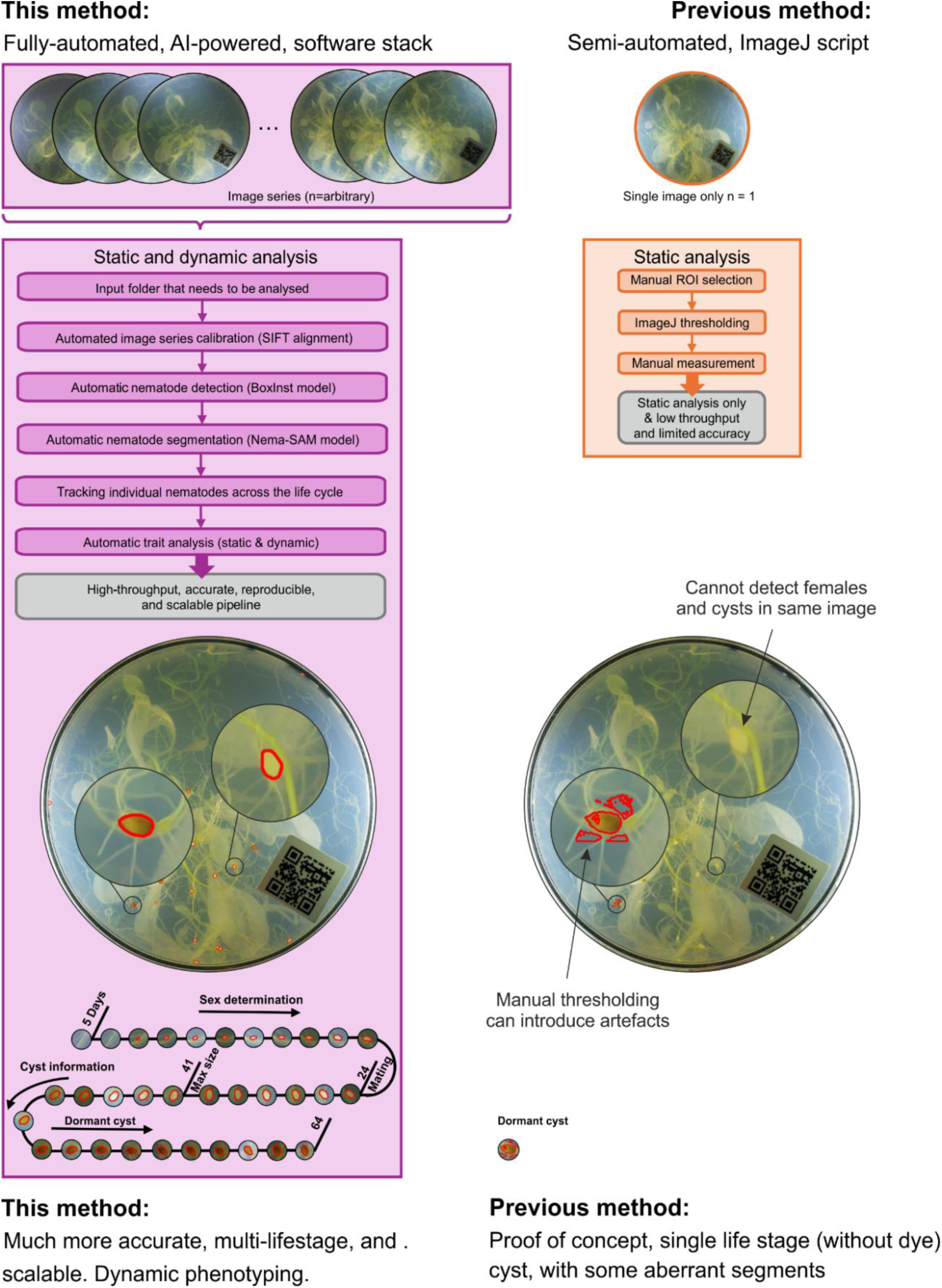
Software comparison between previous method (right) and this method (left). There are no common lines of image analysis code when compared to the previous method (Kranse et al,. 2022). The most striking difference is that the novel method is based on AI, which dramatically improves accuracy, expands the scope of life stages that are measurable, and maintains object permeance over time (thereby enabling for the first time dynamic assessment of individual parasites).

**Figure S5.**
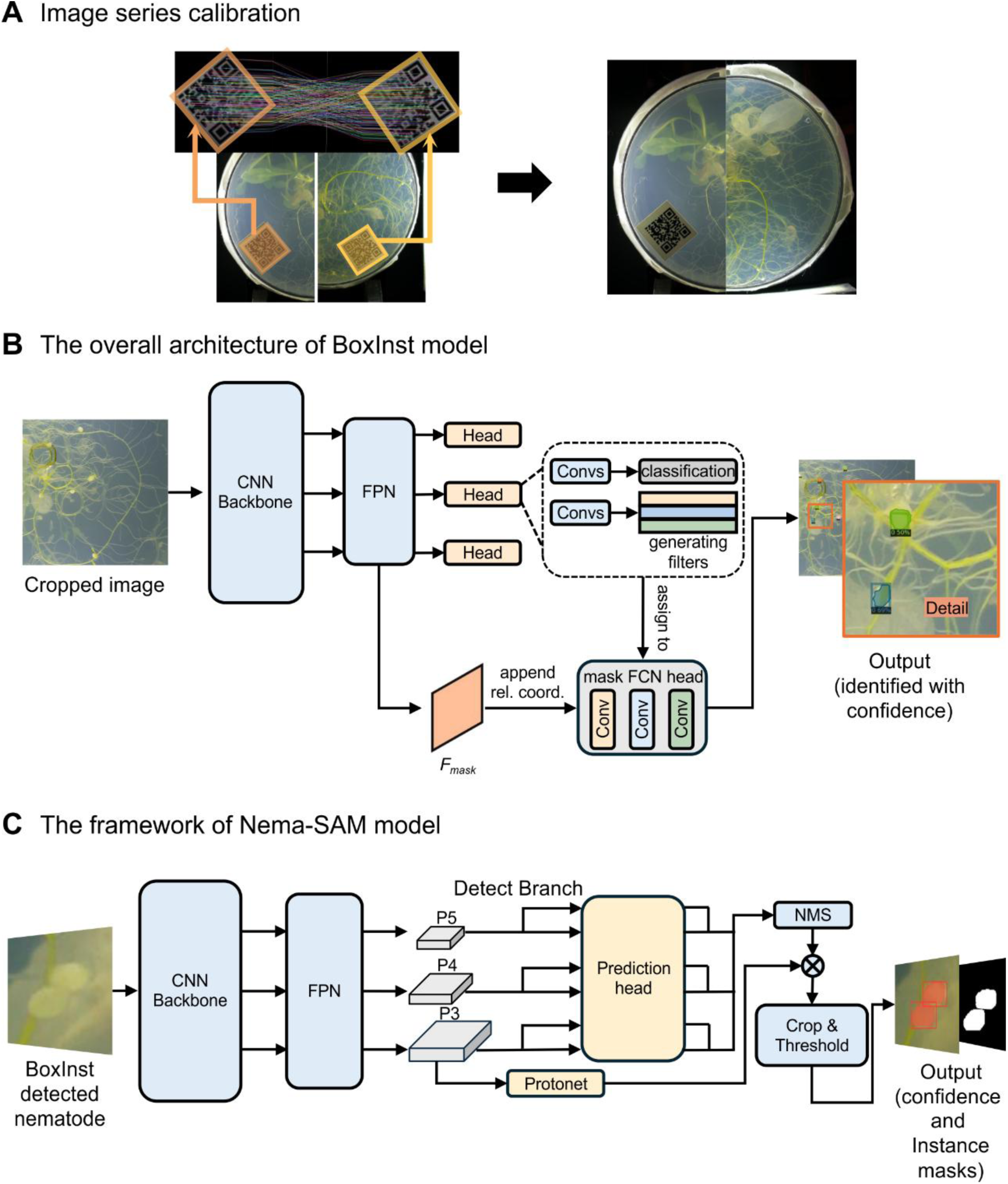
AI architecture. **A)** Image calibration was based on the scale-invariant feature transform (SIFT) algorithm, which detects scale-and rotation-invariant keypoints from embedded QR codes to enable accurate image alignment across timepoints. **B)** Nematode detection was performed using the BoxInst framework, which combined a convolutional neural network (CNN) backbone and feature pyramid network (FPN) to extract multi-scale features. Multiple prediction heads were used for bounding box regression and mask generation, followed by a fully convolutional network (FCN) mask head for each nematode. This stage achieved accurate nematode-level detection, offering spatial localization and initial morphological boundaries, which supported subsequent phenotypic analysis. **C)** Fine-scale morphological phenotypic analysis was performed using the customised Nema-SAM model. It adopted a CNN backbone with a feature pyramid network (FPN) to extract multi-scale features. These features were then passed to a Prototypical Network (Protonet) for object encoding and then to a prediction head for mask generation. Refined instance masks were obtained through non-maximum suppression (NMS), enabling the extraction of phenotypic traits for each individual nematode such as size, length, width, and roundness.

**Figure S6.**
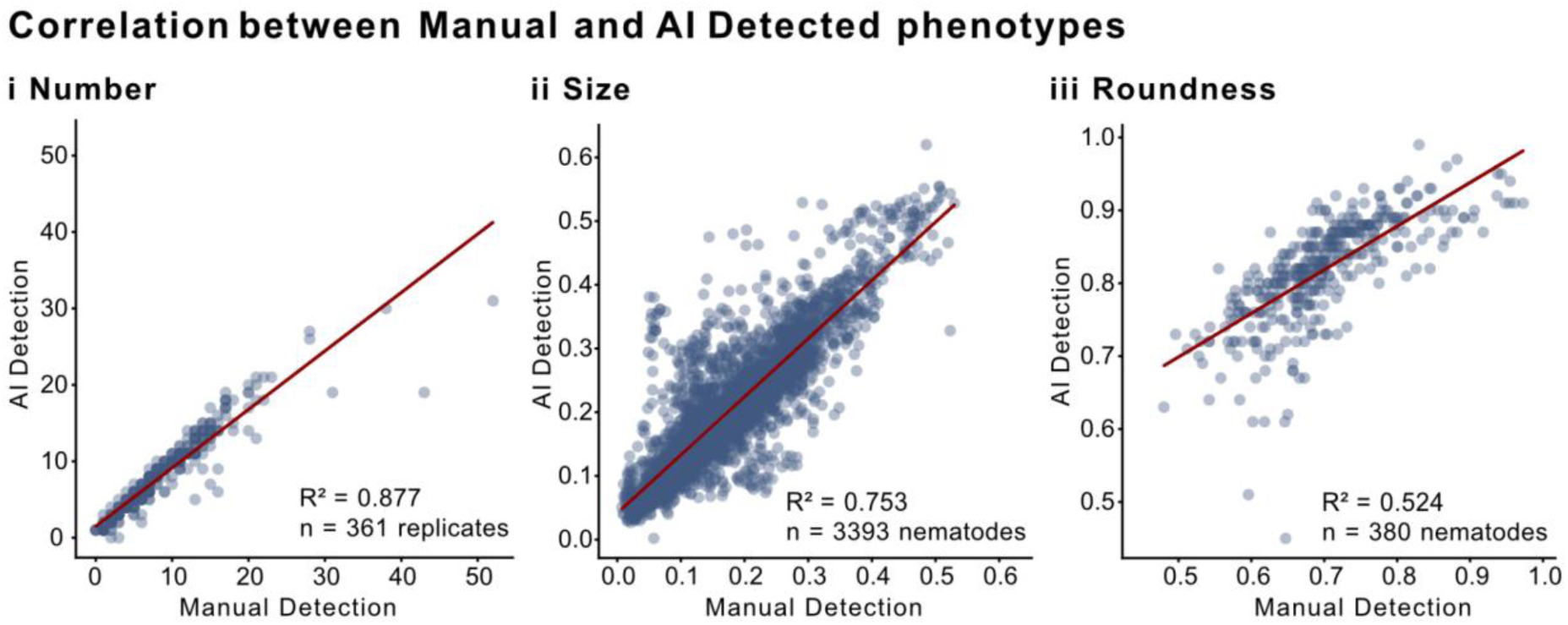
Comparison between manual and automated analyses of nematode phenotypes. Scatter plots compare manual measurements (x-axis) and AI-predictions (y-axis) for three phenotypic traits: (i) number of nematodes per plate (n = 361 replicates, R² = 0.877, P <0.0001), (ii) individual nematode size (n = 3,393, R² = 0.753, P <0.0001), and (iii) individual nematode roundness (n = 380, R² = 0.524, P <0.0001). Each point represents a biological replicate or individual nematode.

**Figure S7.**
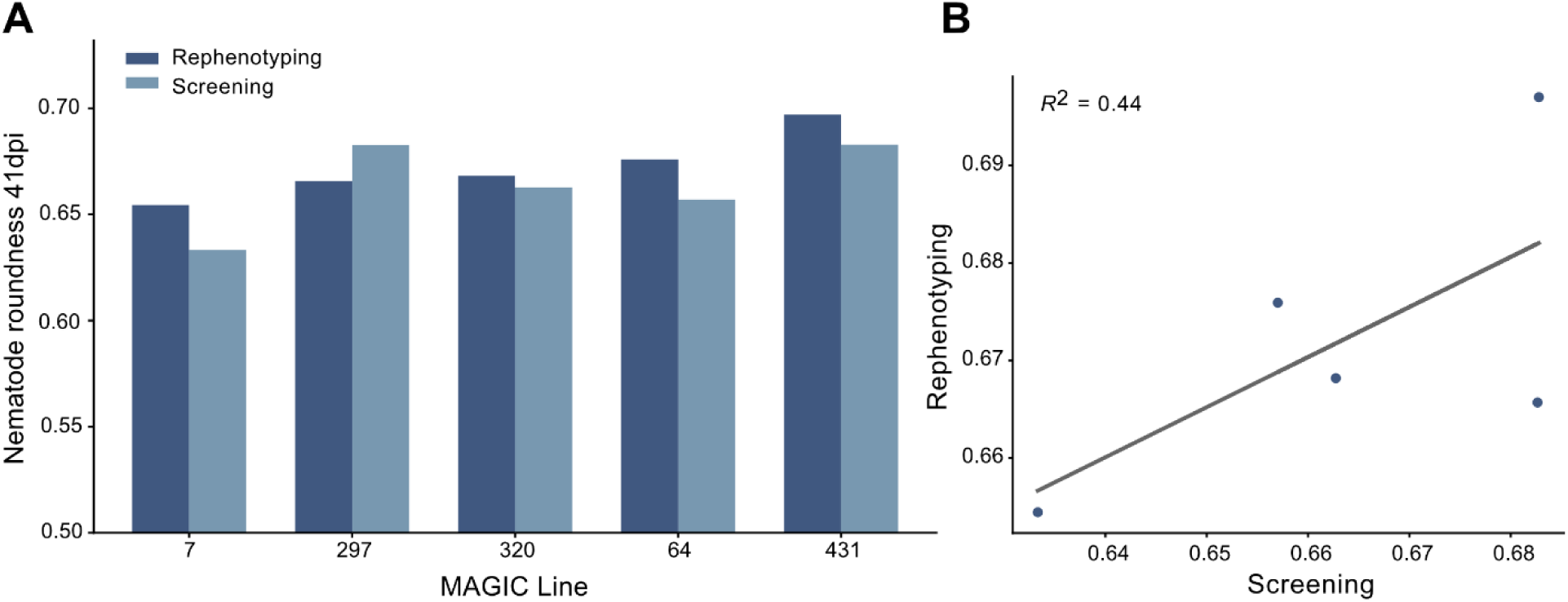
Re-phenotyping of selected MAGIC lines for the roundness phenotype. **A**) Roundness for each of 5 lines from the original screening (light blue) and the repeated re-phenotyping. **B**) X-Y scatter plot comparing roundness of nematodes on the same 5 selected lines from the original screening and the repeated re-phenotyping.

**Figure S8.**
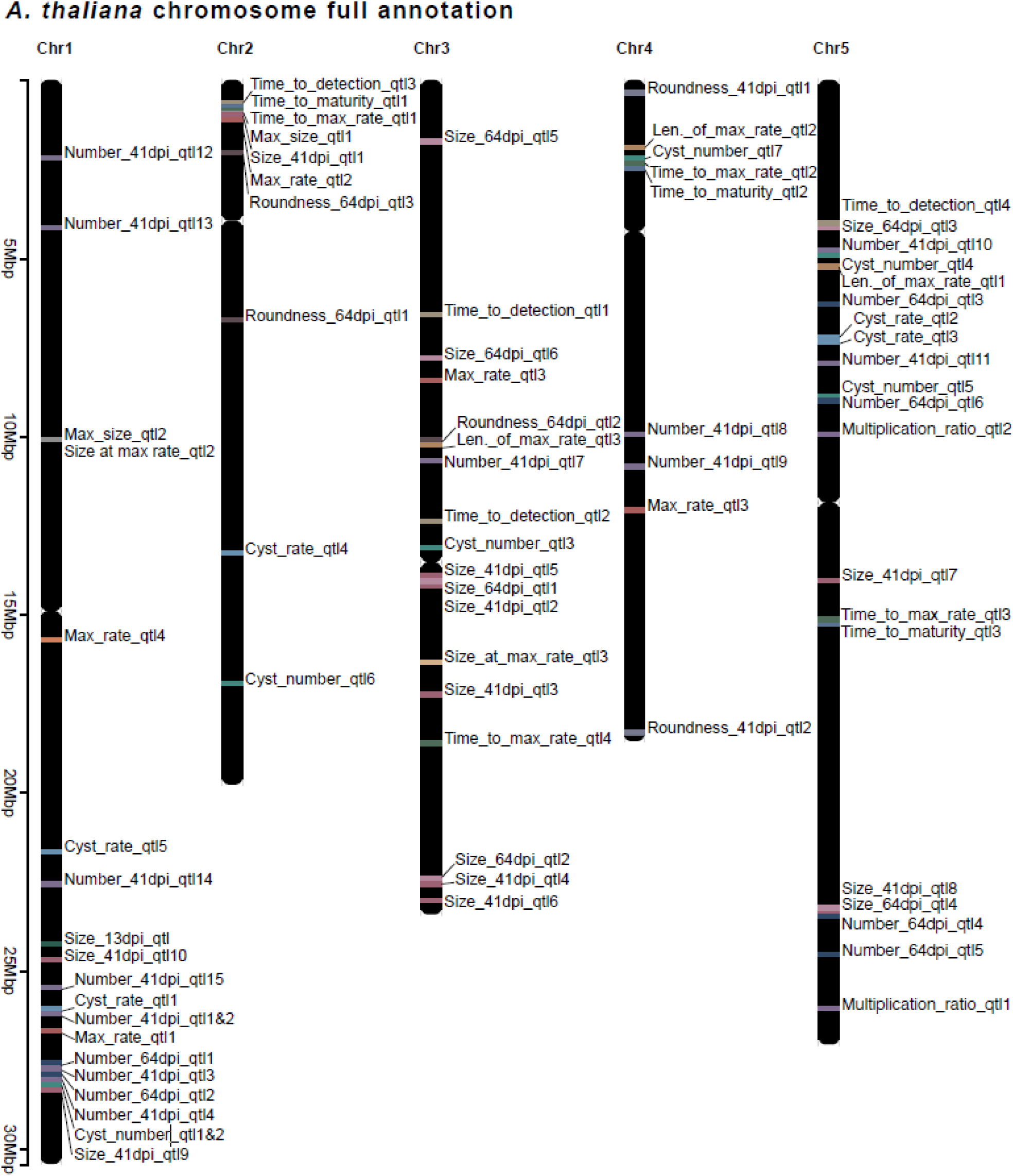
***Arabidopsis thaliana* full annotated chromosome plot.** *Arabidopsis thaliana* chromosome-wide annotation of 78 quantitative trait loci (QTLs) based on summary of all GWAS for all 18 static and dynamic phenotypes. Chromosomal positions are marked along the left axis in Mbp.

**Figure S9.**
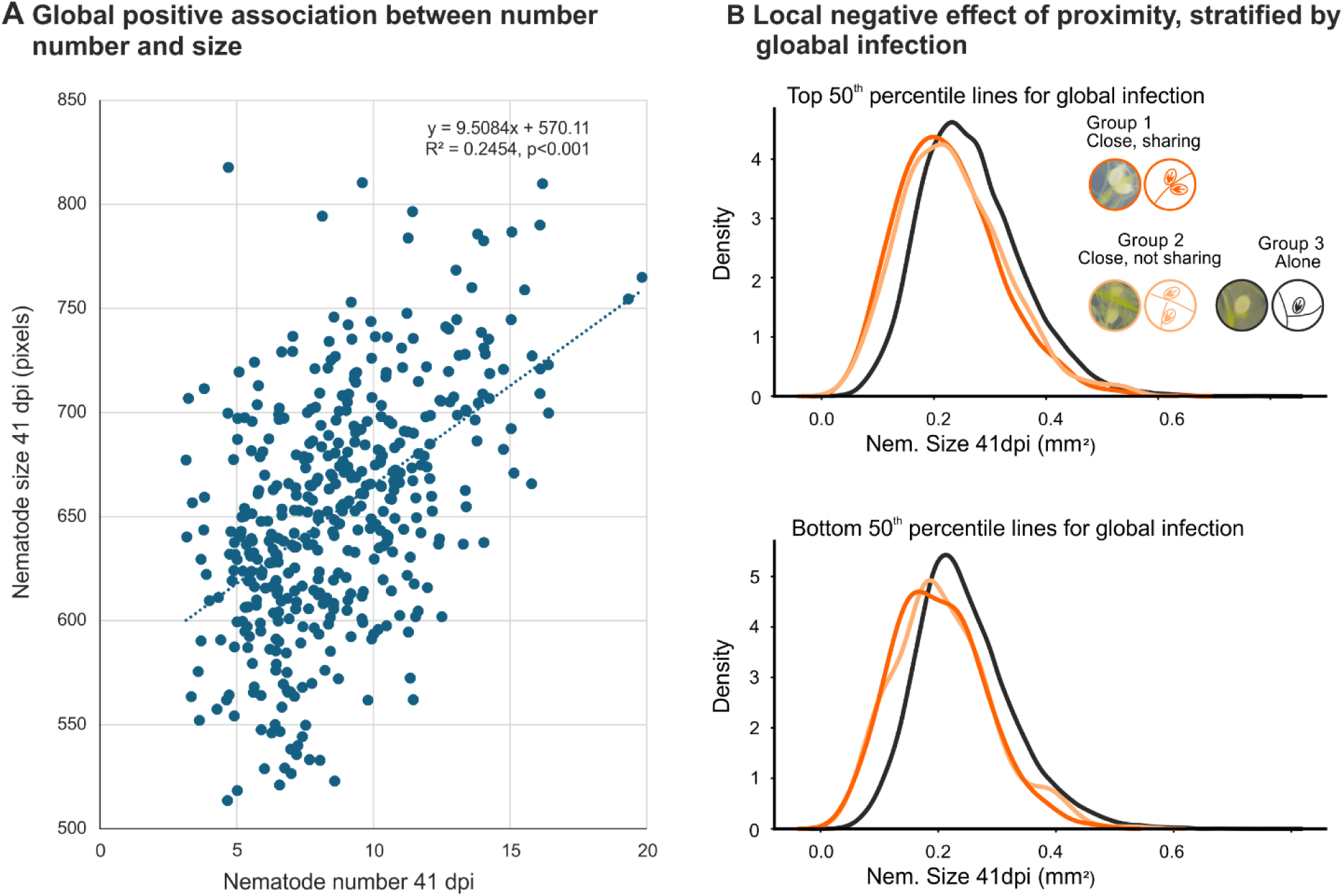
A local negative interaction between nematodes in the context of a global positive association. **A)** scatter plot of nematode number vs nematode size on MAGIC lines at 41 days post infection (dpi) shows overall positive association. **B**) Density plots of local negative effect of nematodes on one another, stratified by number of nematodes per plant (Top 50^th^ percentile shown on top, and the bottom 50^th^ percentile shown on bottom).

**Table S1. Table of all 78 Quantitative Trait Loci (QTL) in the plant associated with 18 nematode phenotypes.**

**Table S2. Summary table of static and dynamic traits (BLUEs) by line.**

